# Unravelling the modulation of intracortical inhibition during motor imagery: An adaptive threshold-hunting study

**DOI:** 10.1101/846931

**Authors:** Cécilia Neige, Dylan Rannaud Monany, Cathy M. Stinear, Winston D. Byblow, Charalambos Papaxanthis, Florent Lebon

## Abstract

Motor imagery (MI) is the mental simulation of an action without any apparent muscular contraction. By means of transcranial magnetic stimulation, few studies revealed a decrease of short-interval intracortical inhibition (SICI) within the primary motor cortex. However, this decrease is ambiguous, as one would expect greater inhibition during MI to prevent overt motor output. The current study investigated the extent of SICI modulation during MI through a methodological and a conceptual reconsideration of i) the importance of parameters to assess SICI (Exp.1) and ii) the inhibitory process within the primary motor cortex as an inherent feature of MI (Exp.2). Participants performed two tasks: 1) rest and 2) imagery of isometric abduction of the right index finger. Using transcranial magnetic stimulation, motor evoked potentials were elicited in the right first dorsal interosseous muscle. An adaptive threshold-hunting paradigm was used, where the stimulus intensity required to maintain a fixed motor evoked potential amplitude was quantified. To test SICI, we conditioned the test stimulus with a conditioning stimulus (CS) of different intensities. Results revealed an Intensity by Task interaction showing that SICI decreased during MI as compared to rest only for the higher CS intensity (Exp.1). At the lowest CS intensities, a Task main effect revealed that SICI increased during MI (Exp.2). SICI modulation during MI depends critically on the CS intensity. By optimising CS intensity, we have shown that SICI circuits may increase during MI, revealing a potential mechanism to prevent the production of a movement while the motor system is activated.

**Highlights:**

- Excitatory and inhibitory neural processes interact during motor imagery, as the motor regions are activated but no movement is produced.
- The current study investigated the extent of short interval intracortical inhibition modulation (SICI) during motor imagery.
- When using optimal settings, SICI increased during motor imagery, likely to prevent the production of an overt movement.

## 1 Introduction

Interactions between excitatory and inhibitory neural processes within the primary motor cortex (M1) are crucial in various cognitive and motor functions (Reis et al., 2008). For example, during motor imagery (MI), the mental simulation of a movement without any apparent muscular contraction, excitatory and inhibitory processes subtly interact as the motor regions are activated but no movement is produced.

Paired-pulse transcranial magnetic stimulation (TMS) protocols provide a quantification of the intracortical processes at the time of the stimulation (Bestmann and Krakauer, 2015). Short-interval intracortical inhibition (SICI) measurements can be obtained by delivering a subthreshold conditioning stimulus (CS), followed 1 to 6 ms later by a second supra-threshold test stimulus (TS) applied through the same coil over M1 (Kujirai et al., 1993). This produces a decrease of motor-evoked potential (MEP) amplitude in comparison to MEP induced by unconditioned TS.

In the conventional paired-pulse TMS paradigm, the peak-to-peak amplitude of the conditioned MEP is expressed as a percentage of the amplitude of the unconditioned MEP, indicating the amount of SICI (Kujirai et al., 1993). This measure depends critically on the CS and TS intensities (Ilić et al., 2002; Peurala et al., 2008; Vucic et al., 2009). First, the TS intensity must be sufficient to recruit the later I-waves suppressed by SICI (Garry and Thomson, 2009; Di Lazzaro et al., 2017). Moreover, changing the CS intensity for a given TS intensity results in a U-shaped SICI curve. In the descending part of this curve, the amount of SICI increases when increasing CS intensity from 50% of the resting motor threshold (rMT), with a peak of inhibition occurring at CS about 80%rMT (Kujirai et al., 1993; Ilić et al., 2002). Then, increasing the CS intensity toward the rMT (i.e., CS intensity >80 rMT (Ilić et al., 2002; Kossev et al., 2003)) leads to the progressive decrease of SICI. This decrease is thought to reflect a “contamination” of the neural process involved in SICI by the recruitment of high-threshold excitatory interneurons. The latter have the potential to override the inhibitory system and are known to contribute to the short interval intracortical facilitation (SICF) phenomenon (Ilić et al., 2002; Kossev et al., 2003; Peurala et al., 2008; Vucic et al., 2009; Wagle-Shukla et al., 2009). Importantly, it must be pointed out that rMT is not a static but rather a state-dependent measure that is subject to the excitability of several cortical and spinal elements excited by the TMS pulse (Groppa et al., 2012; Karabanov et al., 2015). For example, MI decreases the rMT (Facchini et al., 2002; Li, 2007; Grosprêtre et al., 2016) and enhances MEP amplitude (Kasai et al., 1997; Yahagi and Kasai, 1998; Lebon et al., 2012; Grosprêtre et al., 2016) when compared to rest. As suggested by Grosprêtre et al. (2015), these findings bring evidence that cortical cell responsiveness to TMS may increase during MI and this could be mediated, at least in part, by a decrease of inhibitory activity within M1 (Grosprêtre et al., 2016). Indeed, some studies found a reduction of SICI during MI in comparison to rest when using the conventional SICI paradigms (Abbruzzese et al., 1999; Patuzzo et al., 2003; Stinear and Byblow, 2004; Kumru et al., 2008; Liepert and Neveling, 2009). Conversely, other studies failed to observe SICI modulation (Ridding and Rothwell, 1999; Stinear and Byblow, 2004; Sohn et al., 2006; Lebon et al., 2012), indicating that mechanisms underlying SICI modulation during MI remain poorly understood. Notably, the difference between these contradictory results seems to rely on the CS intensity. It appears that only studies fixing the CS intensity at ≥75 rMT found a reduction of SICI during MI.

The aim of the present study was to unravel the SICI modulation observed during MI through a methodological and conceptual reconsideration of: (i) the importance of CS intensity and (ii) the inhibitory process within M1 as an inherent feature of MI. To do so, we designed a pair of experiments in which we varied the CS intensity and determined the TS intensity required to maintain a fixed MEP amplitude for each condition using an adaptive threshold hunting technique (Awiszus et al., 1999; Fisher et al., 2002; Awiszus, 2003; Samusyte et al., 2018; Vucic et al., 2018). This method has been recently developed in order to overcome the potential limitations of conventional paired-pulse TMS protocols, such as large variability in MEP amplitude and a “floor/ceiling effect” when the observed inhibition leads to complete MEP suppression (Cirillo and Byblow, 2016; Cirillo et al., 2018; Van den Bos et al., 2018). The adaptive threshold-hunting technique provides a new opportunity to extend our understanding of physiological mechanisms underlying intracortical inhibition in healthy subjects and it has been recently shown to be more reliable with shorter acquisition time than conventional SICI techniques (Samusyte et al., 2018).

Taking advantage of the adaptive threshold-hunting approach, two different experiments were conducted in order to investigate the evolution of SICI during MI as compared to rest. We used the adaptive threshold-hunting technique in its original form (Fisher et al., 2002) to measure SICI using three CS intensities. We hypothesized a decrease of SICI during MI when compared to rest, as previously observed in the literature with conventional SICI techniques when CS intensity is high (Exp 1). Then, we optimized the adaptive threshold-hunting technique and measure SICI using two low CS intensities. We expected an increase of SICI during MI with low CS when compared to rest, as MI is thought to suppress neural commands at some level of the motor system by inhibitory mechanisms (Jeannerod and Decety, 1995; Jeannerod, 2001).

## 2 Experimental Procedures

### 2.1 Participants

Twenty healthy volunteers (five females; mean age 24.3 years, range 22-27 years; eighteen right-handed as assessed by the Edinburgh Handedness Inventory (Oldfield, 1971)) participated in the current study after providing written informed consent. All volunteers were screened for contraindications to TMS by a medical doctor. The protocol was approved by the University of Burgundy Committee on Human Research and complied with the Declaration of Helsinki. Ten participants were included in Experiment 1 and three of them plus ten other participants were included in Experiment 2.

### 2.2 General procedure

For the two Experiments, participants were comfortably seated on a chair with the forearms supported by a pillow and palms facing down. They were instructed to stay at rest throughout the experiments. For MI trials, participants performed kinesthetic MI of a right tonic index abduction for 3s after an auditory cue. It has been shown that a kinesthetic MI strategy produces a greater muscle-specific and temporally modulated facilitation of the corticospinal pathway, as compared with a visual MI strategy (Stinear et al., 2006). At the beginning of the experiment, participants performed and practiced actual abduction of the right index finger. They then received the following instructions (in French): “When you hear the auditory cue, imagine making an abduction of the right index finger. Try to feel the movement, imagining the muscle contraction and tension that you would expect to experience in actual action. Be sure not to contract any muscles during the task and keep your eyes open” (Lebon et al., 2019).

#### TMS and EMG recordings

Electromyographic (EMG) recordings of the right first dorsal interosseous (FDI) muscle were made with surface Ag/AgCl disposable electrodes in a belly-tendon montage. A ground electrode was placed on the styloid process of the ulna. The EMG signals were amplified and band-pass filtered (10–1000 Hz, Biopac Systems Inc.) and digitized at a sampling rate of 2000 Hz for off-line analysis. Background EMG was monitored for the 100 ms preceding every TMS pulse to ensure a complete muscle relaxation throughout the experiments.

Single-pulse and paired-pulse stimulations were applied with a 70-mm figure-of-eight coil connected to a monophasic Magstim BiStim^2^ stimulator (The Magstim Co., Whitland, UK). The coil was placed over the left M1, tangentially to the scalp with the handle pointing backward and laterally at 45° away from the midsagittal line, resulting in a posterior-anterior current flow within M1. The optimal stimulation site on the scalp (hotspot) was defined as the location eliciting the largest MEP amplitude in the FDI and was marked on the scalp. The conventional rMT was determined as the lowest stimulation intensity required to evoke at least 5 MEPs of 50 μV out of 10 stimulations (Rossini et al., 1999) and then was used to set CS intensities.

#### Adaptive threshold-hunting technique

In Experiment 1, based on the corresponding literature, the hunting-threshold was defined as the TS intensity (expressed in percentage of the maximal stimulator output (%MSO)) required to elicit a MEP_target_ in the relaxed FDI of 0.2 mV in peak-to-peak amplitude. This 0.2 mV MEP amplitude was chosen in accordance to numerous previous studies using the adaptive threshold-hunting technique (Fisher et al., 2002; Awiszus, 2003; Vucic et al., 2006; Menon et al., 2015; Cirillo and Byblow, 2016; Cirillo et al., 2018; Samusyte et al., 2018; Van den Bos et al., 2018). This 0.2mV fixed MEP value has been shown to lie on the middle of the steepest portion of the stimulus response curve plotted on a logarithmic scale (Fisher et al., 2002; Vucic et al., 2006, 2018). In the current study, the TS intensity required to elicit the MEP_target_ at rest corresponded on average to 109% rMT (range 103-118 % rMT), which is similar to a previous study (Cirillo and Byblow, 2016). The adaptive threshold-tracking single-pulse TMS technique was used to first compare the unconditioned TS intensity (%MSO) required to maintain this fixed MEP_target_ amplitude at rest vs. during MI.

The adaptive threshold-hunting paired-pulse TMS technique was then used to investigate SICI at rest and during MI. To elicit SICI at rest and during MI in Experiment 1, we delivered high CS intensities: 60%, 70% and 80% of the rMT. The TS intensity was adjusted to reach the MEP_target_. A 2 ms interstimulus interval between CS and TS was chosen based on a similar previous study investigating SICI modulation during motor imagery (Lebon et al., 2012).

In Experiment 2, the hunting threshold was defined as the TS intensity (%MSO) required to elicit a MEP in the relaxed FDI of at least 50% of MEP_max_ in peak-to-peak amplitude. This MEP_target_ amplitude in Experiment 2 has been chosen since a TS delivered at a low intensity (i.e., below 110% rMT, as it was the case in Experiment 1), could fail to evoke late indirect waves, and limit SICI magnitude (Garry and Thomson, 2009). This subject-specific relative MEP value is half of the individual’s maximum MEP amplitude value. MEP_max_ was calculated with a stimulus/response curve performed at the beginning of the experiment. We recorded eight MEPs for each stimulus intensity starting at 110% of rMT with incrementing steps of 10% rMT up to MEP_max_ (Kukke et al., 2014; Pitcher et al., 2015). In the current study, the TS Intensity required to elicit the MEP_target_ at rest (mean MEP amplitude: 1.4 mV ± 0.81) corresponded on average to 124% rMT (range 112-150 %rMT). In the same way as for Experiment 1, the adaptive threshold-hunting single-pulse TMS technique was used to compare the unconditioned TS intensity (%MSO) needed to elicit the MEP_target_ at rest vs. during MI.

To elicit SICI at rest and during MI in Experiment 2, we delivered low-intensity CS, i.e. 50% and 60% rMT. The CS was delivered 2 ms prior to TS. TS intensity was adjusted in each condition.

For both experiments, the order of the experimental conditions was randomized across participants. An available online freeware (TMS Motor Threshold Assessment Tool, MTAT 2.0), based on a maximum-likelihood Parameter Estimation by Sequential Testing (PEST) strategy was used with “assessment without a priori information” in line with previous studies (Cirillo and Byblow, 2016; Cirillo et al., 2018). The stimulation sequence always began with the TS at 37 %MSO. One experimenter held the coil over M1, while the other indicated whether or not the MEP amplitude was ≥0.2 mV (Experiment 1) or ≥ 50%MEPmax (Experiment 2). The predictive algorithm then determined the next TS intensity to be delivered and was stopped after thirty stimulations, which provides sufficient accuracy for the threshold estimate according to previous studies (Awiszus, 2003, 2014; Ah Sen et al., 2017).

## 3 Data analysis

First, the unconditioned TS intensity required to elicit the MEP_target_, at rest and during MI was quantified and expressed in %MSO in both experiments.

Then, to probe the influence of the different CS on TS intensity, the amount of SICI (expressed in INH%) was quantified for each condition using the following equation (Fisher et al., 2002):

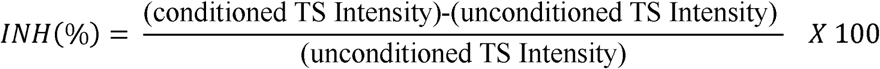

where positive values indicate inhibition and negative values indicate facilitation.

Background root mean square (RMS) of the surface EMG was calculated during the 100 ms epoch prior to TMS to ensure the absence of muscle contraction in each condition.

## 4 Statistical analysis

Statistical analyses were performed using Statistical Program for the Social Sciences (SPSS) version 24 software (SPSS Inc., Chicago, IL, USA). Normality of the data distributions was verified using the Shapiro-Wilk test. Homogeneity of variances was assessed by Mauchly’s test and a Greenhouse-Geiser correction was applied if the sphericity assumption was violated. Pre-planned post-hoc analyses were performed on significant interactions after applying a Bonferroni correction for multiple comparisons. Corrected p values for multiple comparisons are reported in the results section. The α level for all analyses was fixed at .05. Partial eta squared (ηp^2^) values are reported when results are statistically significant to express the portion of the total variance attributable to the tested factor or interaction. Values in parentheses in the text represent mean ± SD.

First, a Student’s two-tailed paired sample *t-*test was used to compare the unconditioned TS Intensity (%MSO) between Rest and MI for both experiments. Then, an analysis of variance (ANOVA) was performed on SICI measurements (INH %) with two within-subject factors: Task_2_ (Rest vs. MI) and CS Intensity_3_ (CS 60% rMT vs. CS 70% rMT vs. CS 80% rMT) for Experiment 1.

For Experiment 2, ANOVA was performed on SICI measurements (INH %) with two within-subject factors: Task_2_ (Rest vs. MI) and CS Intensity_2_ (CS 50% rMT vs. CS 60% rMT). Additional analyses were performed to control for potential methodological biases. The RMS values were compared across conditions in both experiments, using the same analyses described above.

## 5 Results

### 5.1 Methodological considerations

The analysis of the pre-trigger background EMG level for the unconditioned TS (Table 1) yielded no significant difference between rest and MI neither in Experiment 1 (*t*(9) = 1.133; p=.287) nor in Experiment 2 (*t*(12) = -1.017; p=.329). In addition, the ANOVAs for RMS values for conditioned TS revealed no significant main effects or interactions for Experiment 1 (all p > .07) and Experiment 2 (all p > .14). Therefore, any changes observed in the subsequent measurements cannot be attributed to differences in EMG levels prior to the TMS pulse.

**Table 1:**
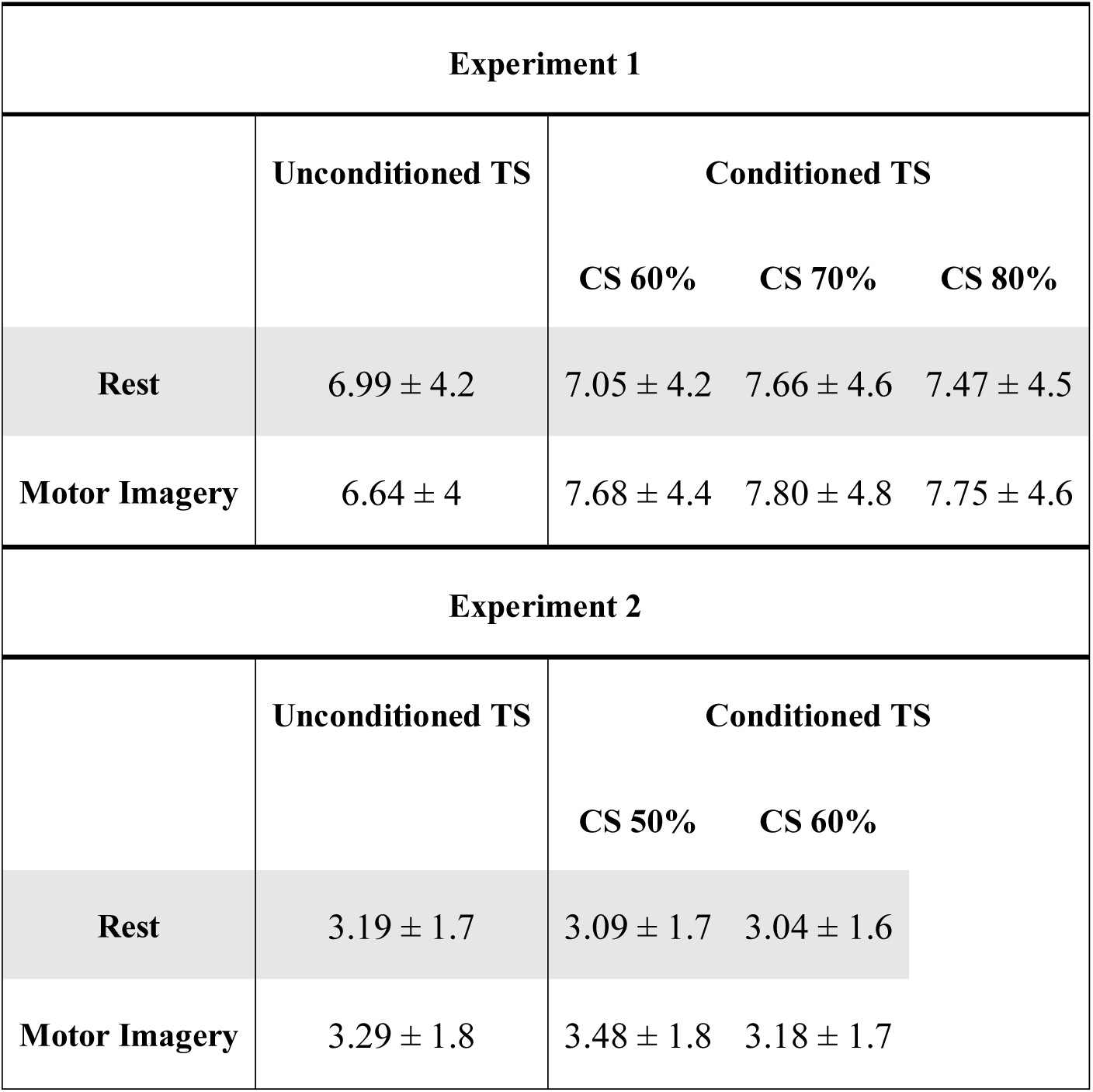
Pre-trigger root mean squared EMG expressed in μV (mean ± SD).

| Experiment 1 |  |  |  |  |
| --- | --- | --- | --- | --- |
|  | Unconditioned TS | Conditioned TS |  |  |
|  |  | CS 60% | CS 70% | CS 80% |
| Rest | $6.99 \pm 4.2$ | $7.05 \pm 4.2$ | $7.66 \pm 4.6$ | $7.47 \pm 4.5$ |
| Motor Imagery | $6.64 \pm 4$ | $7.68 \pm 4.4$ | $7.80 \pm 4.8$ | $7.75 \pm 4.6$ |
| Experiment 2 |  |  |  |  |
|  | Unconditioned TS | Conditioned TS |  |  |
|  |  | CS 50% | CS 60% |  |
| Rest | $3.19 \pm 1.7$ | $3.09 \pm 1.7$ | $3.04 \pm 1.6$ | |
| Motor Imagery | $3.29 \pm 1.8$ | $3.48 \pm 1.8$ | $3.18 \pm 1.7$ | |

### 5.2 Unconditioned TS Intensity

Figure 1 illustrates the unconditioned TS Intensity obtained at rest and during MI in Experiment 1 (left panel) and Experiment 2 (right panel) in both groups of participants. Two-tailed paired sample *t-*tests revealed that the unconditioned TS Intensity required to reach the MEP_target_ was significantly lower during MI than at rest in both Experiment 1 (49.3 ± 11.3 vs. 53.3 ± 10.6; *t*(9) = 3.381; p=.008) and Experiment 2 (43.4 ± 8.7 vs. 45.5 ± 13.6; *t*(12) = 2.976; p=.012). This result indicates that MI leads to increase corticospinal excitability. Note that unconditioned TS Intensity at rest was higher in Experiment 1 in comparison to Experiment 2 (*t*(21) = 3.364; p=.003), due to greater rMT for individuals recruited in the first experiment (Experiment 1, mean rMT = 48.6 %MSO (range 27-59); Experiment 2, mean rMT = 36.7 %MSO (range 28-47).

**Figure 1:**
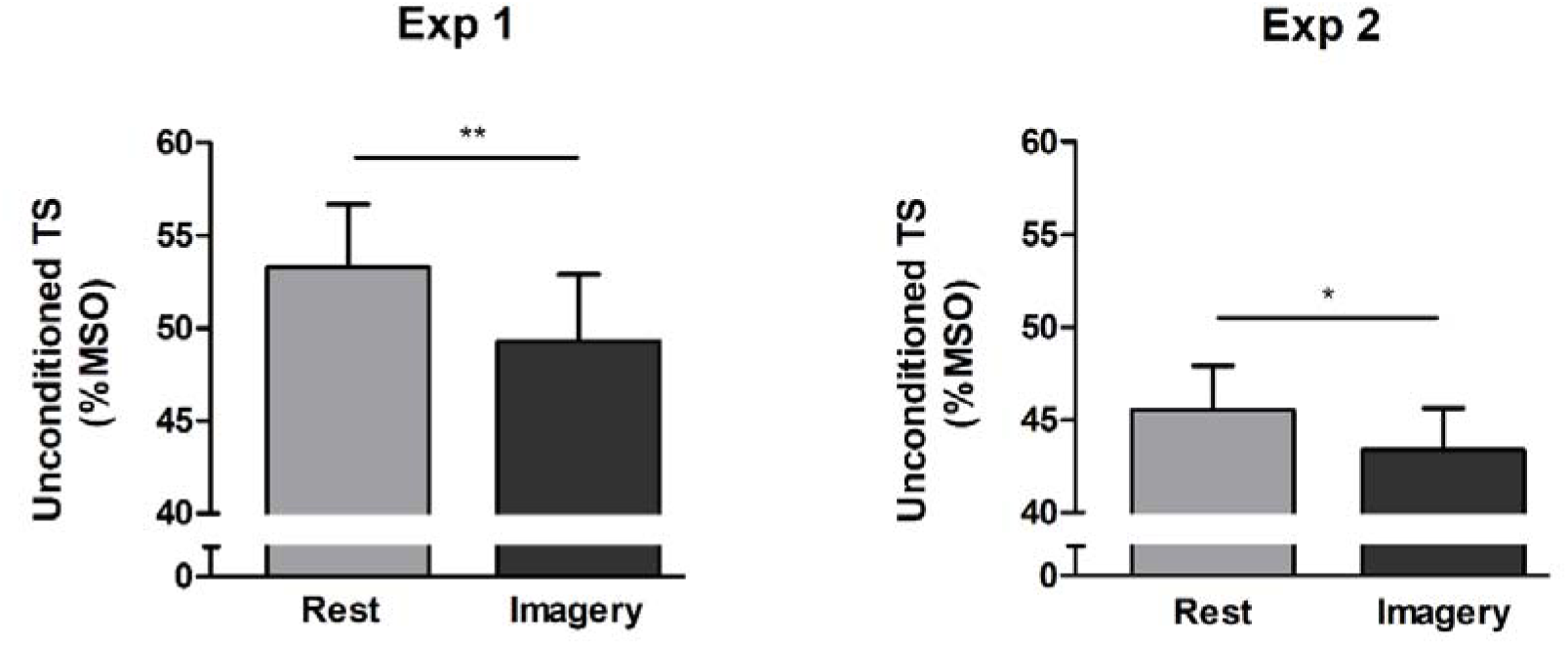
Mean ± SE for the Unconditioned TS Intensity obtained with the hunting-threshold technique at rest and during motor imagery in the two Experiments. *p < .05; **p<.01

### 5.3 Conditioned TS Intensity (SICI)

#### Experiment 1

Figure 2 illustrates the percentage of inhibition (SICI) obtained at rest and during MI for the three CS intensities. ANOVA revealed no main effects of Task (F_(1,9)_ < 1, p = .582) or CS intensity (F_(2,18)_ < 1, p = .871). However, there was a Task by CS intensity interaction (F_(2,18)_ = 9.086, p = .002; ηp^2^ = .502). Post-hoc analyses revealed that there was less SICI during MI than at rest only for the CS intensity of 80% rMT (p = .004; Figure 3 for typical raw MEP recordings).

**Figure 2:**
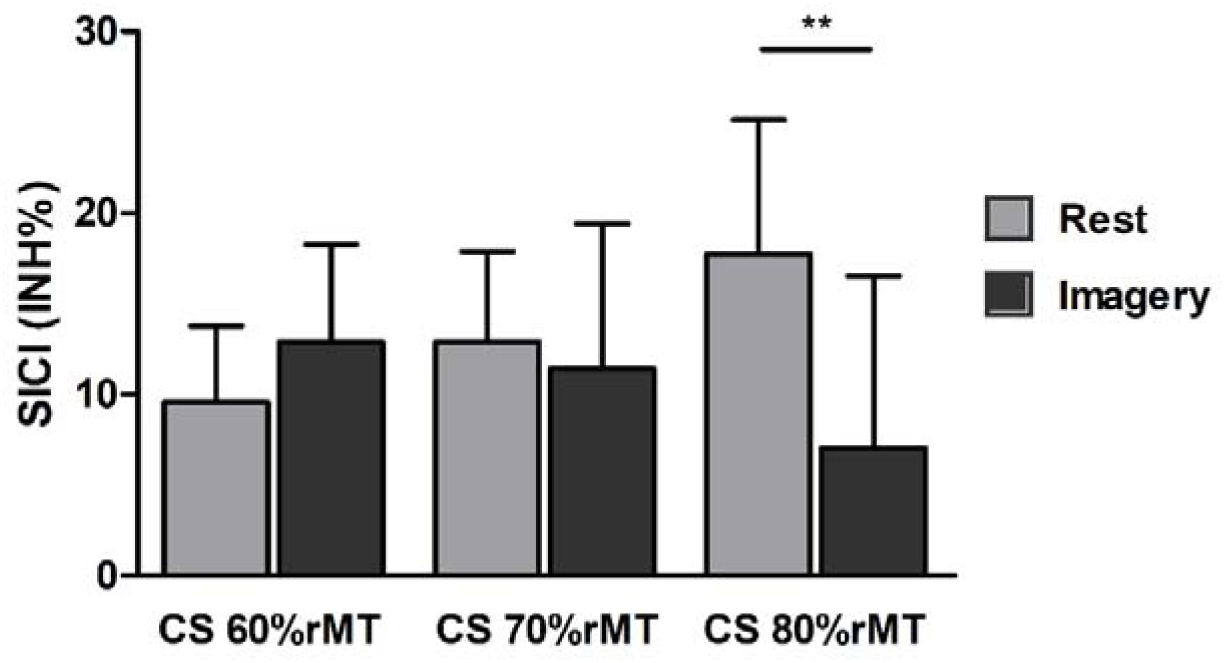
Mean ± SE for SICI obtained in Experiment 1 with the hunting-threshold technique and according to the three conditioning stimulus (CS) intensities expressed in percentage of the rMT at rest and during motor imagery. **p<.01

**Figure 3:**
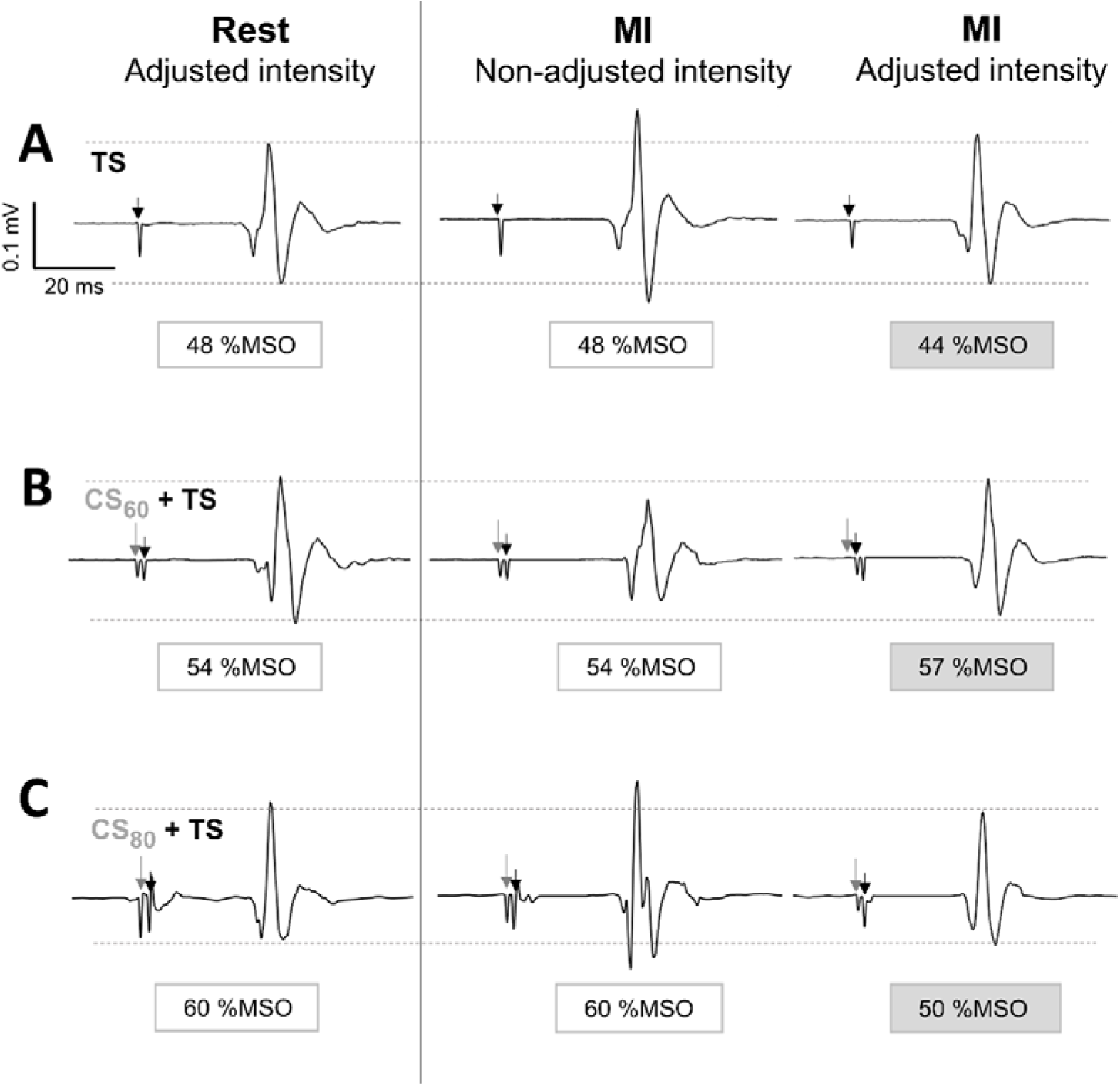
Typical raw MEP recordings of the right FDI muscle in Experiment 1. **(A)** Unconditioned Test Stimulus (TS, black arrow). At rest, the TS intensity was adjusted to elicit the MEP_target_ (0.2 mV). During motor imagery (MI), the same TS intensity elicited a greater MEP amplitude; it was therefore decreased to evoke the MEP_target_. **(B)** Paired-pulse protocol where the conditioning stimulus (CS, grey arrow) was delivered at an intensity of 60% of resting motor threshold (CS60). In comparison to rest, TS intensity during MI was increased to evoke the MEP_target_. (C) Paired-pulse protocol where the CS was delivered at an intensity of 80%rMT (CS80). In comparison to rest, TS intensity during MI was decreased to evoke the MEP_target_.

#### Experiment 2

Figure 4 illustrates SICI results (INH%) obtained at rest and during MI for two different CS intensities. ANOVA revealed a main effect of Task showing that SICI is higher during MI when compared to rest (+35.4 ± 9.3 INH% vs. +30.5 ± 10.3 INH%, F_(1,12)_ = 4,838; p = .048; ηp^2^ = .287). However, no significant main effect of CS intensity (F_(1,12)_ = 2,543, p = .137) nor Task by CS Intensity interaction (F_(1,12)_ < 1, p = .467) was found.

**Figure 4:**
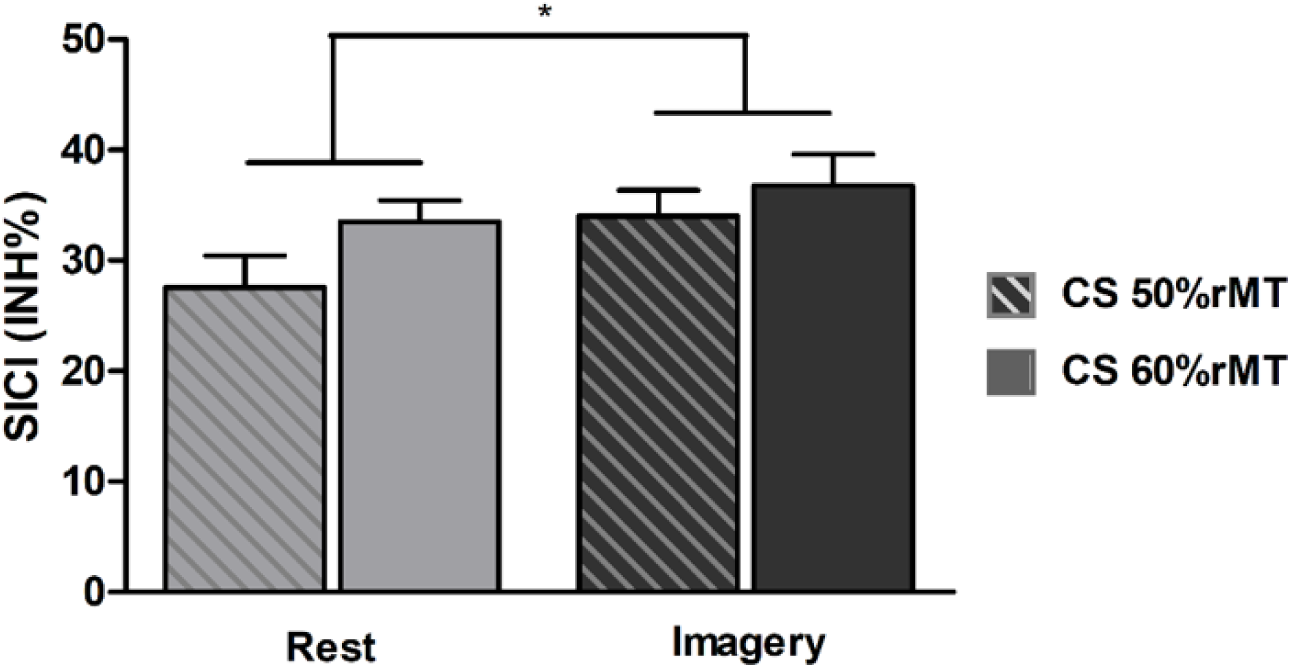
Mean ± SE for SICI obtained in Experiment 2 with the hunting-threshold technique and according to the two conditioning stimulus (CS) intensities expressed in percentage of the rMT at rest (grey) and during motor imagery (black). *p < .05

## 6 Discussion

The main objective of the present study was to investigate the modulation of SICI during MI, with a particular focus on the effect of the CS intensity. For the first time, we reported that the CS intensity chosen to evaluate SICI led to opposite conclusions regarding SICI modulation during MI. When tested with a high CS intensity there is less SICI during MI, as compared to rest. On the contrary, when tested with optimizing low CS intensities there is a greater SICI during MI than at rest. These findings are the first to strength the idea that MI requires motor command inhibition by active processes acting at a cortical level.

### Corticospinal excitability increases during MI

While the major objective of this study was to focus on SICI, we also analyzed the level of corticospinal excitability (unconditioned TS intensity) at rest and during MI for the two experiments. By using the adaptive hunting threshold single-pulse TMS technique, we found that the minimum TS intensity required to elicit the MEP_target_ was lower during MI when compared to rest. This result replicates and extends earlier findings showing that corticospinal excitability is higher during MI (Karabanov et al., 2015; Grosprêtre et al., 2016; Ruffino et al., 2017), regardless of the MEP_target_ amplitude.

### SICI is lower during motor imagery when tested with high CS intensity

In Experiment 1, SICI decreased during MI as compared to rest only for the highest CS intensity (CS intensity set at 80% rMT), as previously observed in the literature with conventional SICI techniques. This decrease has been suggested to explain the corticospinal excitability increase observed during MI (Abbruzzese et al., 1999; Patuzzo et al., 2003; Stinear and Byblow, 2004; Kumru et al., 2008; Liepert and Neveling, 2009).

However, there is now compelling evidence that cortical cell responsiveness to TMS increased during MI, as observed by a decrease of the rMT in comparison to rest (Grosprêtre et al., 2016). Therefore, for a similar subthreshold CS intensity based on the rMT calculated when the subject is at rest, this same CS intensity for the MI condition is closer to the motor threshold. The investigations of SICI at rest have found that increasing the CS intensity leads to the recruitment of high-threshold excitatory interneurons that may contaminate SICI, motivating us to carry out the current study. The result obtained here does indeed show that SICI was lower during MI when compared to rest only for high CS intensity (80% rMT). We may therefore suggest that this decrease in SICI could be the result of using CS intensities that were too high and that produced an unwanted recruitment of excitatory interneurons. These findings lead us to reconsider the modulation of SICI underlying MI, taking into account the selection of CS intensity.

### SICI is higher during motor imagery when tested with low CS intensities

In Experiment 2, we found for the first time that SICI was greater during MI (vs. at rest) when using low CS intensities (i.e. 50% rMT and 60% rMT). With optimal TMS settings, we revealed an important component of neural processes within M1 during MI.

Neuroimaging studies have shown that brain networks underlying MI and actual movement execution extensively overlap, supporting the elaboration of motor commands during MI (Hétu et al., 2013). During MI, however, motor commands may be stopped at some level of the motor system by active inhibitory mechanisms to prevent them from being sent to peripheral effectors (Jeannerod and Decety, 1995). It has been hypothesized that the neural activation within motor and pre-motor areas during MI is blocked by inhibitory mechanisms preventing the overt action (Jeannerod and Decety, 1995; Jeannerod, 2001). Because intracortical networks within M1 could be considered as the final cortical modulators of motor output (Cowie et al., 2016), a possible explanation for these findings is that the increased SICI would prevent the production of an overt movement when the mental representation of that movement is activated. These findings strengthen the idea that MI requires motor command inhibition by active processes acting at a cortical level.

One could argue that the SICI increase, i.e. more inhibition, cannot be at play simultaneously to the corticospinal excitability increase, i.e. more facilitation, in the specific effector involved during MI. However, it is important to keep in mind that MEP amplitude results from the balance between inhibitory and excitatory processes along the corticospinal tract including both cortical and spinal-segmental contributions. Moreover, neuroimaging studies provide evidence that MI is also supported by a network involving motor and premotor regions including cortical and subcortical structures (Hétu et al., 2013). Therefore, corticospinal facilitation could possibly originate from these regions, outside M1 and exert their influence via direct or indirect pathways (Reis et al., 2008). By contrast, the modulation of SICI observed in this study could reflect the crucial role played by cortical interneurons within M1 in the fine-tuning neural processes required during MI.

### MEP_target_ amplitude considerations

In the current study, different MEP_target_ amplitudes were chosen for the two experiments and this deserves discussion. In Experiment 1, based on the existing literature, a fixed MEP amplitude value of at least 0.2 mV was tracked for all participants (Fisher et al., 2002; Vucic et al., 2018), corresponding to an average TS intensity of 109% rMT (range 103-118 %rMT), a relatively low intensity. It is known that SICI predominately inhibits the late I-waves (I2, I3 and I4) and that a TS delivered at a low TMS intensity (i.e., below 110% rMT) could fail to evoke late I-waves and thus limits the detection of SICI (Garry and Thomson, 2009; Di Lazzaro et al., 2017).

In order to answer our second hypothesis and to optimize the conditions of SICI measure, a subject-specific MEP amplitude of at least 50% of MEP_max_ was tracked for all participants in Experiment 2, corresponding on average to 124 % rMT (range 112-150 %rMT), a moderate TS intensity known to generate the greatest measure of (Garry and Thomson, 2009; Wagle-Shukla et al., 2009; Amandusson et al., 2017; Van den Bos et al., 2018). Reliable stimulus-response curves can be acquired in less than 4 minutes (van de Ruit et al., 2019) and allow personalisation of the hunting MEP_target_ amplitude. Therefore, future studies using the adaptive threshold hunting technique to investigate intracortical mechanisms could consider target 50% MEP_max_.

### Perspective

The results obtained in the current study have demonstrated that modulation of SICI during MI depends critically on CS intensity. Importantly, other cognitive conditions that share analogous control mechanisms and neural circuits with overt movements, such as motor preparation or action observation, are known to selectively modulate corticospinal excitability and to affect SICI (Naish et al., 2014; Duque et al., 2017). Supporting this view, a recent study measured SICI using a range of CS intensities at rest and during a warned simple reaction time task (Ibáñez et al., 2019). The results showed that show that SICI changes that occurred during the task could be either larger or smaller than at rest depending on the intensity of the CS. Together, these findings also confirmed that testing SICI using a wide range of CS intensities provides a more nuanced interpretation of possible GABAergic changes in M1 than testing with a single CS intensity (Ibáñez et al., 2019). We believe that the adaptive threshold hunting paradigm could be useful in further studies to assess SICI during various cognitive and motor states.

### Conclusion

Overall, this study provides initial evidence that the intensity of the CS crucially affects SICI measurement during MI when compared to rest. The previously reported decrease in SICI during MI could be due to inappropriate TMS settings, with high CS intensities leading to the unwanted recruitment of excitatory interneurons. With low CS intensities, we show that SICI is greater during MI than at rest, probably to prevent the production of an overt movement when the mental representation of that movement is activated. Future studies should consider optimizing the SICI stimulation protocol by careful adjustment of the CS intensity.

## Declaration/conflicts of interest

No conflicts of interest, financial or otherwise, are declared by the authors.

## Funding

This work was supported by the French “Investissements d’Avenir” program, project ISITE-BFC (contract ANR-15-IDEX-0003).

## Notes

### Competing Interest Statement

The authors have declared no competing interest.

